# Evolution of virulence of a plant RNA virus in age-diverse host populations

**DOI:** 10.64898/2026.03.24.713855

**Authors:** José L. Carrasco, Christina Toft, Santiago F. Elena

## Abstract

Natural host populations are intrinsically age-structured, and developmental stages differ in susceptibility and within-host dynamics, meaning age compositions can impose distinct selective pressure on pathogens. However, how host age structure shapes viral evolution remains largely untested. We experimentally evolved turnip mosaic virus (TuMV) for five passages in *Arabidopsis thaliana* populations spanning seven demographic regimes (from juvenile-dominated to mature-dominated cohorts). We quantified disease progression, symptom severity and viral load, cross-inoculated evolved lineages across host stages to build quantitative infection matrices, and performed whole-population sequencing at passages 1 and 5 to infer selection coefficients and test for parallelism. Disease traits changed strongly with passage, demography, and their interaction. Disease progression evolved faster in older host populations, whereas severity showed no detectable dependence on median age, implying demographic reweighting of virulence components. Viral load increased through passages and positively correlated with severity, linking within-host fitness to symptoms. Cross-assays revealed a non-nested but modular infection network: juvenile-evolved lineages specialized on pre-bolting plants, whereas lineages from intermediate/older demographies were more stage-generalist. Genomically, both parallel and demography-specific solutions emerged, with a dense cluster of recurrent changes in VPg and several synonymous variants showing consistent or sign-flipping selection across demographies. In conclusion, host age structure is a primary ecological driver of virulence evolution, determining whether viruses evolve faster disease timing versus stronger severity, and whether they specialize or generalize across host stages. These results integrate phenotypes with genome-level responses and suggest that manipulating crop age pyramids could steer virus evolution toward less damaging outcomes.

---

Host homogeneity is rare in natural populations, meaning that pathogens typically encounter host populations whose individuals are genetically diverse and differ in their susceptibility to infection (Pfenning et al. 2001; González et al. 2019; Elena et al. 2025). Interactions between pathogens and hosts that vary genetically in susceptibility have been well studied (Chang et al. 1992; Schmid-Hempel & Koella 1994; Brown & Tellier 2011; Hillung et al. 2014; Navarro et al. 2022), as has the influence of developmental stage on host resistance (Chang et al. 1992; Heath 1993; Rupe & Gbur 1995; Izhar & Ben-Ami 2015; Melero et al. 2023). Variation in disease susceptibility across ages or developmental stages is a key component of host heterogeneity, with the potential to affect host-pathogen outcomes and epidemiological dynamics (Ben-Ami 2019). These differences may imply that a virus can be adapted to a particular host developmental stage yet fail to infect others effectively. In nature, interactions between host developmental stage and susceptibility have been observed in many animal and plant systems across a wide range of pathogens. In animals, young individuals are generally more susceptible to bacterial and viral infections than older ones (Krakowka & Koestner 1976; Klinge et al. 2009; Cleton et al. 2014). However, for some pathogens such as fungi, older animals may show increased susceptibility (Bradley et al. 2019). In plants, hosts commonly become more resistant to bacterial and fungal pathogens as they mature (García-Ruiz & Murphy 2001; Kus et al. 2002; Shibata et al. 2010), a pattern associated with age-related resistance (ARR) (Hu & Yang 2019). ARR, however, is not universal and depends on the specific pathosystem: some plants become more susceptible to particular fungal pathogens (Miller 1983) and RNA viruses (Huang et al. 2020; Melero et al. 2023) as they develop.

Plants finely coordinate growth and development to optimize fitness by mounting rapid, appropriate responses to diverse stresses. The life cycle of flowering plants can be viewed as a succession of distinct phases, including the transition from a juvenile vegetative stage to a mature reproductive stage. The transition to flowering is governed by a complex genetic network that integrates endogenous and environmental cues (Amasino 2010). Although flowering itself is not the developmental transition required for increased pathogen resistance (Wilson et al. 2013), several studies have described links between flowering and defense. Depending on the pathogen and host genotype, viral infection can delay reproduction (Pagán et al. 2008; Shukla et al. 2018) or accelerate flowering (Korves & Bergelson 2003; Winter et al. 2011). These patterns are consistent with resource-allocation theory: plants have a limited pool of resources that must be distributed among functions throughout the life cycle, creating trade-offs (Weiner 2004; Boege & Marquis 2005). As plants develop, they eventually shift resources from biomass accumulation and structural growth to reproduction. A similar reallocation occurs during pathogen attack, when plants must deploy resources strategically to overcome infection. A particularly well-studied example is the trade-off between defense and floral development, especially consequential for species that reproduce only once in their lifetime. Moreover, pathogens encounter different host defense responses depending on plant developmental stage (Kus et al. 2002; Korves & Bergelson 2003). These stage-specific differences can elicit different pathogen strategies to counter host defenses. Developmental stage-specific states, such as shifts in reactive oxygen species or reprogramming of hormone crosstalk pathways (Khan et al. 2014; Mhamdi & Breusegem 2018; Melero et al. 2024), influence host susceptibility and thus present changing scenarios for pathogens.

Despite the attention given to developmental stage during infection, we still lack information on how different host stages impose distinct selective environments for viruses, thereby modulating viral evolution. Most eco-evolutionary theory assumes a link between virulence and transmission (Anderson & May 1982), positing a compromise between infection duration (*i.e*., the time during which a pathogen can propagate and infect other hosts) and the damage inflicted on the host. Notably, the virulence-transmission trade-off appears to be affected by host’s life history (Ebert & Herre 1996; Perlman 2008; Izhar & Ben-Ami 2015). Although parasite-induced host mortality is the most commonly used measure of virulence for horizontally transmitted parasites (Alizon et al. 2009), some pathogens employ alternative strategies that ensure reproduction and transmission without prematurely killing the host. Because development entails substantial reallocation of host resources (Harper & Ogden 1970), viruses infecting the same host genotype may nonetheless face different selective pressures depending on host age. Consequently, adaptive mutations that enhance viral fitness at one developmental stage may be neutral, or even selected against, at another.

Here, we sought to better understand how variation in host developmental stage within populations influences virus evolution. We used populations of *Arabidopsis thaliana* that differed in demographic composition to evaluate the evolutionary dynamics of their natural parasite, turnip mosaic virus (TuMV; species *Potyvirus rapae*, genus *Potyvirus*, family *Potyviridae*). Experimental plant populations differed in the proportion of individuals at three developmental stages: (*i*) juvenile vegetative, characterized by resource allocation to growth; (*ii*) bolting, which marks the developmental transition (Pouteau & Albertini 2009); and (*iii*) flowering, during which mature plants allocate resources to reproduction. These stages have already been shown to affect the severity of TuMV infection (Melero et al. 2023) and to involve age-specific reprogramming of defense responses (Melero et al. 2024). We evaluated how median host population age influences the evolution of TuMV virulence, viral genetic diversity, identified candidate beneficial mutations, and tested whether adaptation was host age-specific.

## Methods

### Plants and viruses

*Arabidopsis thaliana* (L.) HEYHN, ecotype Col-0, was used in this study. Plants were grown in a BSL2 climatic chamber under an 16 h light/8 h dark photoperiod (LED tubes, PAR 90 - 100 μmol·m⁻^2^·s⁻^1^), at 24 °C during light and 20 °C during dark, with 40% relative humidity. The substrate consisted of 50% DSM WNR1 R73454 (Kekkilä Professional, Vantaa, Finland), 25% grade 3 vermiculite, and 25% perlite (3 - 6 mm). Biological pest control was implemented using *Stratiolaelaps scimitus* and *Steinernema feltiae* (Koppert Co., Málaga, Spain).

The infectious plasmid p35STunos, containing a cDNA of the YC5 TuMV isolate from calla lily (*Zantedeschia* sp.; GenBank accession AF530055.2) under the cauliflower mosaic virus 35S promoter and *nos* terminator (Chen et al., 2003), was used to infect *Nicotiana benthamiana* DOMIN.

For inoculation of *A. thaliana*, infected *N. benthamiana* leaf tissue was ground and suspended at 100 mg/mL in inoculation buffer (50 mM phosphate buffer, pH 7, 3% PEG 6000), supplemented with 10% Carborundum suspension (100 mg/mL in the same buffer). Two leaves of 3-week-old plants were mechanically inoculated with 5 μL of this suspension. Mock-inoculated plants received buffer plus Carborundum only.

### Experimental evolution design

Ancestral TuMV was experimentally evolved in *A. thaliana* in populations with different age compositions through five serial passages. For passage *t* = 1, plants were inoculated with sap from infected *N. benthamiana*. For subsequent passages, symptomatic plants were pooled at 14 dpi and used as inoculum for the next passage.

Host populations consisted of varying proportions of plants at three developmental stages (Figure 1): pre-bolting (juvenile, ordinal age category 1), bolting (transition to reproductive growth, ordinal age category 2), and post-bolting (mature, ordinal age category 3). Four independent lineages were evolved under demographic conditions (pre-bolting:bolting:post-bolting): expansive 12:8:4 (median ordinal age category Ã = 1.5), stationary 4:16:4 (Ã = 2) and 8:8:8 (Ã = 2), and constrictive 4:8:12 (Ã = 2.5). For homogeneous populations (expansive 24:0:0, Ã = 1; stationary 0:24:0, Ã = 2; and constrictive 0:0:24, Ã = 3), three new lineages were evolved plus three previously reported by Melero et al. (2023). Each population included 24 plants.

**Figure 1.**
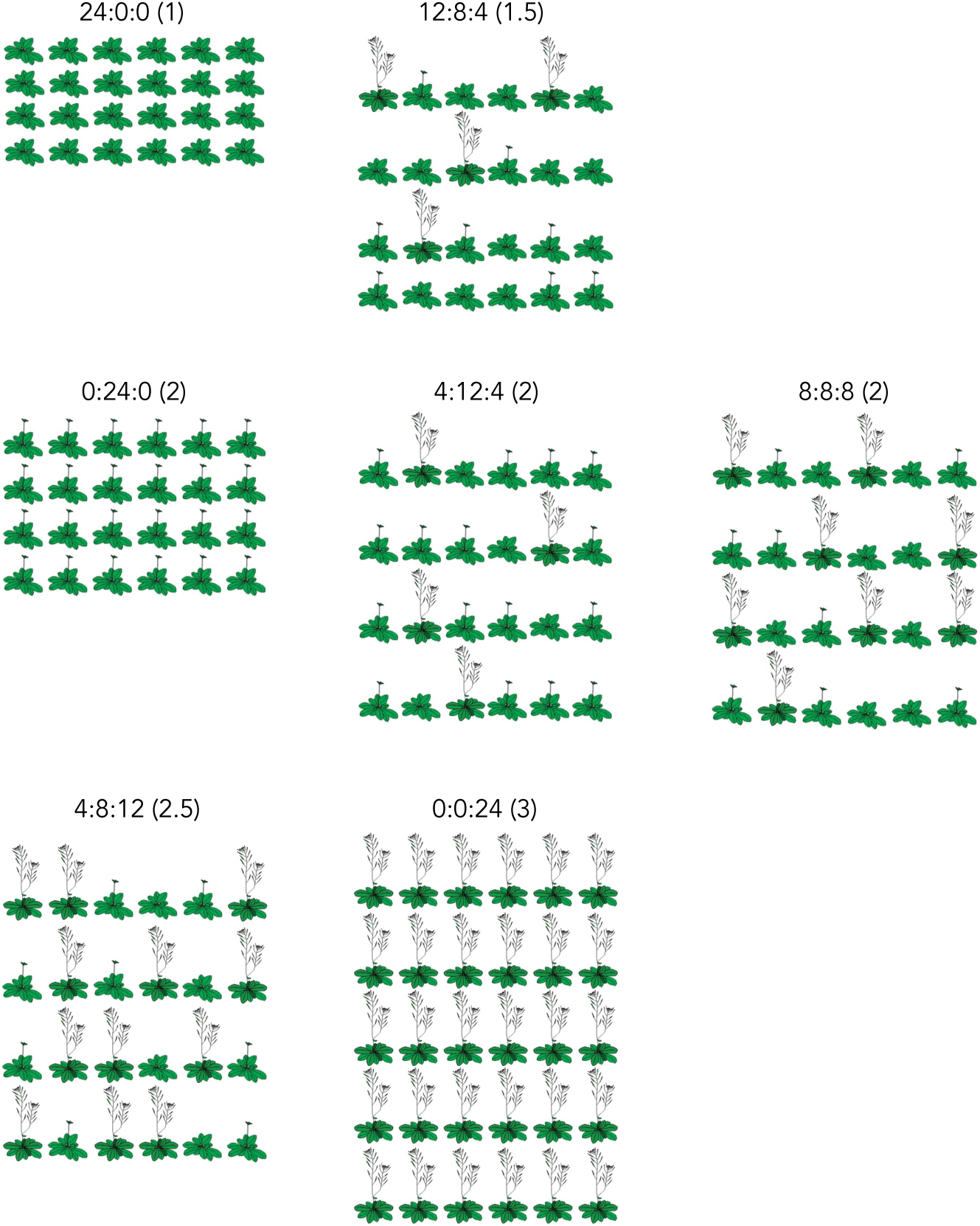
Schematic of the seven host population age compositions used for experimental evolution. Numbers in parenthesis show the median ordinal age categories (Ã) for each population. The upper row shows “expansive” populations dominated by juveniles (Ã = 1 or 1.5), the middle row shows “stationary” populations with balanced age classes (Ã = 2 in all three cases), and the lower row shows “constrictive” populations dominated by older plants (Ã = 2.5 or 3). Each population comprised 24 *A. thaliana* plants allocated to pre-bolting (juvenile), bolting (transition), or post-bolting (mature) stages in the indicated proportions.

### Phenotyping disease

Symptoms were scored daily for each plant using a semi-quantitative scale (0 = no symptoms, 5 = systemic necrosis; Butković et al. 2021), except for passage 2. These data were used to estimate: (*i*) incubation period (mean time to first symptoms) via Kaplan-Meier regression, (*ii*) mean symptoms severity over 14 days, (*iii*) maximum symptoms severity at 14 dpi, (*iv*) area under the disease progress stairs (AUDPS; Simko & Piepho 2012), and (*v*) are under symptom intensity progress stairs (AUSIPS; Kone et al., 2017). AUDPS and AUSIPS were computed using the R package ‘agricolae’ version 1.3-2 (de Mendiburu 2020).

### RNA extraction, estimation of viral load, and preparation of samples for RNA-seq

Viral accumulation was quantified after passages *t* = 1 and *t* = 5. Symptomatic plants were pooled, frozen and homogenized under liquid nitrogen. RNA was extracted using the NZY Plant/Fungi RNA isolation kit (NZYtech, Lisbon, Portugal) following manufacturer’s recommendations.

Viral load was measured by absolute real-time quantitative RT-PCR (RT-qPCR) using a TuMV standard curve and the primers TuMV F117 forward (5’-CAATACGTGCGAGAGAAGCACAC-3’) and F118 reverse (5’- TAACCCCTTAACGCCAAGTAAG-3’), amplifying a 173 bp fragment of the CP cistron (Corrêa et al., 2020). Briefly, standard curve consisted of eight serial dilutions of the *in vitro* synthesized TuMV genome prepared in total plant RNA purified from healthy *A. thaliana* plants.

Reactions were run in 10 μL using qPCRBIO SyGreen 1-step Go Hi-ROX System (PCRBiosystems, London, UK) on an ABI StepOne Plus Real-time PCR System (Applied Biosystems, Foster City CA, USA). Cycling: RT 45 °C (10 min), denaturation at 95 °C (2 min), then 40 cycles of 95 °C (5 s) and 60 °C (25 s), followed by melting curve analysis. Negative controls included RNA from healthy plants and water. Each sample was run in triplicate. Results were analyzed using the StepOne software 2.2.2 (Applied Biosystems).

Library preparation and Illumina sequencing were performed by Novogene (Cambridge, UK) using a NovaSeq X Plus Series (PE150) platform and an Lnc-stranded mRNA-seq library method, rRNA depletion, directional library preparation, 150 bp paired-end, and 6 Gb raw data per sample. Novogene did a quality check of the libraries using a Qubit 4 Fluorometer (Thermo Fisher Scientific, Waltham MA, USA), qPCR for quantification, and bioanalyzer for size distribution detection.

### Infection matrix

Evolved lineages were used to inoculate *A. thaliana* plants at the three aforementioned developmental stages. Infection progress was monitored daily basis; viral load was quantified 14 dpi. Trait values and log-viral load were transformed into integers (0 - 9) (Moury et al. 2021) for nestedness and modularity analysis using R packages ‘bipartite’ version 2.18 and ‘igraph’ version 1.5.0.9008. Nestedness was estimated with WNODF (Almeida-Neto & Ulrich 2011) and modularity with the spinglass algorithm (Newman & Girvan 2004). Significance was tested against 10,000 null matrices generated under the B model using Patefield (1981) algorithm.

### Bioinformatics methods

Fastq quality was assessed with FASTQC (Andrews 2010) and MultiQC (Ewels et al., 2016). Reads were trimmed with BBDuk (https://sourceforge.net/projects/bbmap/) to remove adapters, first 10 nucleotides and low-quality bases (*Q* < 10), discarding reads < 80 nucleotides long. Parameter: *ktrim* = r, *k* = 31, *mink* = 11, *qtrim* = r, *trimq* = 10, *maq* = 5, *forcetrimleft* = 10, and *minlength* = 80. Reads were mapped to the TuMV YC5 genome using BWA-MEM (Li, 2013). SAM files were processed with SAMtools (Danecek et al. 2021), and duplicates marked with GATK MarkDuplicates version 4.6.1.0 (McKenna et al. 2010).

TuMV variant calling was performed with LoFreq (Wilm et al. 2012), retaining SNVs with allele frequency *p* > 0.05 and coverage > 100 reads (Kofler et al. 2011; Spitzer et al. 2020). Per site heterozygosity (*h*) was computed using custom R scripts. It was analyzed using the aligned rank transform (ART) method for nonparametric factorial ANOVA, implemented in the R package ‘ARTool’ version 0.11.2.

### Inferences of selection coefficients and tests of parallelism

For each genomic position × treatment × replicate, allele counts were aggregated for the ancestral (*t* = 1) and final (*t* = 5) time points. Alternative-allele frequency at site *i* and at each timepoint *t*, *p_i_*(*t*), was defined as the sum of the all non-reference allele counts divided by the total coverage on site *i*. Thus, *p_i_*(1) and *p_i_*(5) refer to alternative-allele frequences at the two sampled time points, respectively, and allele-frequency changes were calculated as Δ*p_i_* = *p_i_*(5) − *p_i_*(1).

To quantify consistent allele-frequency changes across replicates, we applied the Cochran-Mantel-Haenszel (CMH) test (Kofler et al. 2011) to each genomic position within the same demographic conditions (Cuevas et al. 2002; Hillung et al. 2014; Longdon et al. 2018; Bertels et al. 2019; Navarro et al. 2022). For each site, 2×2 contingency tables (alternative *vs* reference at *t* = 5 *vs t* = 1) were constructed for each replicate, and CMH statistics and *P*-values were computed across replicates (Kofler et al. 2011; Schlötterer et al. 2015; Spitzer et al. 2020). Direction of allele-frequency change was taken from the CMH output, which is derived from the sign of the odds ratio (OR). CMH *P*-values were corrected for multiple testing using Benjamini-Hochberg false-discovery-rate (FDR) procedure. A site was classified as exhibiting parallel evolution within a demographic condition if it met three criteria; (*i*) an FDR-corrected CMH adjusted *P* < 0.05; (*ii*) at least two-thirds of replicates showing allele-frequences change in the same direction; and (*iii*) an absolute mean Δ*p_i_* ≥ 0.02.

Finally, selection rate constants for dominant allele at locus *i* (*s_i_*) were estimated for each site and replicate using the logit-slope approximation *s_i_* ≍ [logit *p_i_*(5) − logit *p_i_*(1)]/Δ*t* (Ewens, 2004). Sampling variances were estimated using the delta method applied to Binomial variances under read depth at *t* = 1 and *t* = 5. Replicate-specific estimates were combined using inverse-variance meta-analysis to obtain treatment-level 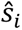 (Paris et al. 2019; He et al. 2020). This approach is closely related to estimators used in other approaches but adapted for two timepoint data [*e.g*., WFABC (Foll et al. 2014), HMM-based diffusion methods (Bollback et al. 2008; Mathieson & McVean, 2013) and CLUES2 (Vaughn & Nielsen 2024)].

### General statistical analyses

The evolution of daily symptom severity was analyzed using a generalized linear mix model (GLMM). In this model, dpi was treated as a repeated-measures factor, evolutionary passage and demography were included as orthogonal between-subjects fixed factors, and independent lineages were modeled as a random factor nested within demographic condition. Tests of within-subject effects were corrected for violations of sphericity using the Greenhouse-Geisser adjustment.

For the traits mean symptom severity, maximum symptom severity at 14 dpi, AUDPS, AUSIPS, and viral load, we fitted a generalized linear model (GLM) with evolutionary passage and demographic condition as fixed orthogonal factors, and lineage as a random factor nested within demographic condition. In addition to univariate analyses, a multivariate analysis of variance (MANOVA) was performed on the four disease-related traits using Wilks’ Λ statistic and the same model structure. To explore trait correlations, we conducted a principal component analysis (PCA) with Varimax rotation and Kaiser normalization. Component importance was assessed based on eigenvalues and their contribution to explained variance.

To estimate rates of phenotypic evolution for each disease-related trait and viral load, time-series data were modeled using ARIMA(*p*, *d*, *q*) models following Elena et al. (2025). Model selection was performed with the R package ‘forecast’ version 8.23.0, using automatic estimation of the Box-Cox transformation parameter λ.

In GLM and GLMM, the 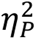 statistic was used to quantify effect sizes (how much of the variance in a dependent variable is explained by a specific factor, after accounting for other factors in the model). Conventionally, 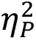 < 0.05 are taken as small effects, 0.05 ≤ 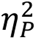 < 0.15 as medium effects and 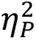 ≥ 0.15 as large effects.

All graphs were generated using the ‘ggplot2’ package version 3.4.0 (Wickham, 2009). All these analyses were conducted in R version 4.5.2 (R Core Team, 2025) within RStudio version 2025.09.2+418 (Posit Software, Boston MA, USA) or SPSS version 30.0.0.0 (IBM Corp., Armonk NY, USA).

## Results

### The evolution of disease progression and symptom severity depends on host population age composition

Across five serial passages, all disease-related traits showed pronounced changes whose magnitude and direction depended on the host population age composition. Univariate GLM/GLMM analyses for each trait (incubation period, AUDPS, AUSIPS, mean symptom severity, and maximum symptom severity at 14 dpi) consistently detected significant effects of passage, host population age composition, and their interaction, with mostly large effect sizes (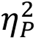), as detailed in the legends of Supplementary Figs. S1 - S5. For example, incubation period and symptoms traits each exhibited highly significant between-subjects effects of passage and host population age composition, as well as significant passage × host population age composition interactions, indicating that the tempo and intensity of disease evolution depended on the host age structure experienced during evolution.

A MANOVA on the five disease traits corroborated these patterns, revealing highly significant and large-magnitude effects of passage, host population age composition, and their interaction (Supplementary Table S1; Wilks’ Λ, all *P* < 0.001, 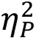 ≥ 0.425).

Traits were not independent. Controlling for Ã, AUDPS was negatively correlated with the incubation period (*r_p_* = −0.999, 4 d.f., *P* < 0.001), whereas AUSIPS correlated positively with mean symptom severity (*r_p_* = 1, 4 d.f., *P* < 0.001) and with final symptom severity at 14 dpi (*r_p_* = 0.940, 4 d.f., *P* = 0.005).

Given MANOVA detected strong multivariate effects of passage, host population age composition, and their interaction across the set of disease traits, the outcomes are best interpreted in a multivariate space rather than trait-by-trait. Moreover, the observed correlations among traits indicates redundancy and potential collinearity; therefore, we used PCA to reduce dimensionality and extract orthogonal composite axes that capture shared variance while enabling clearer inference and visualization with fewer multiple comparisons. This analysis summarized disease phenotypes into two axes (Fig. 2A - 2B). PC1 (63.1% variance) primarily captured symptom severity [large positive loadings for AUSIPS (0.928), mean severity (0.948), and severity at 14 dpi (0.955); near-zero weight for incubation period (–0.112) and AUDPS (0.119)], whereas PC2 (35.1%) represented disease progression timing, loading strongly and oppositely on incubation period (–0.991) and AUDPS (0.345). Thus, PC1 reflects how severe disease becomes, and PC2 how fast disease develops, largely independent of severity (Fig. 2A - 2B).

**Figure 2.**
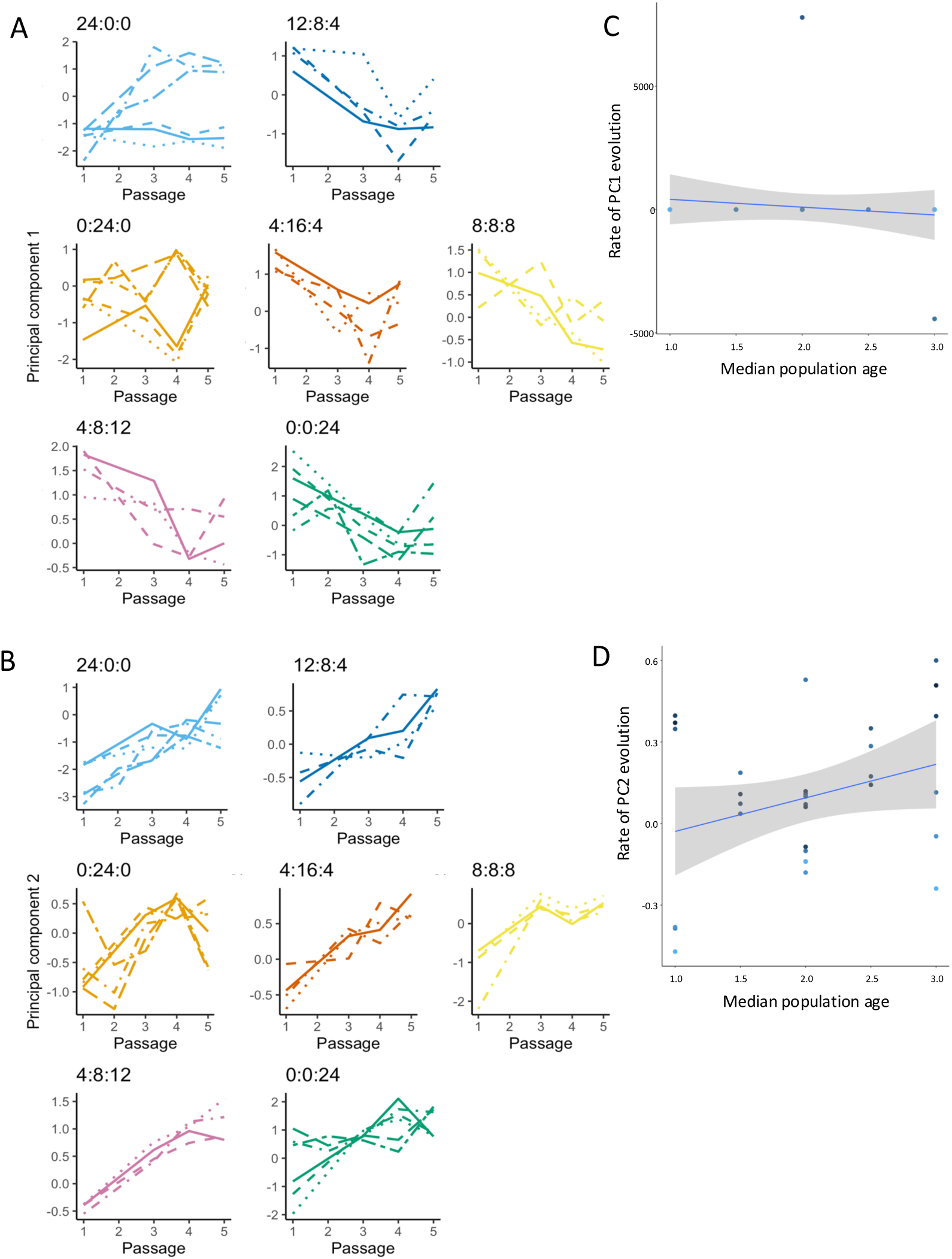
Principal components of disease phenotype and their evolutionary dynamics. (A) Evolution of PC1 (symptom severity) across passages under each host population age composition. (B) Evolution of PC2 (disease progression timing). Different evolutionary lineages are distinguished by line type; host age compositions are indicated above panels. (C) Dependence of the rate of evolution for PC1 and (D) PC2 on median of the ordinal age categories, Ã. Solid lines show linear fits (PC1: *R*^2^ = 0.000, *F*_1, 24_ = 0.000, *P* = 1.000; PC2: *R*^2^ = 0.065, *F*_1, 24_ = 1.084, *P* = 0.309); grey bands indicate ±1 SD around the regression line.

### Within-host viral fitness in infected individuals depends on the host age pyramid structure

Viral load per infected host (RT-qPCR) increased during experimental evolution but did so heterogeneously across host population age compositions (Fig. 3A). A GLM including passage and demographic condition (with lineage as random, nested within host population age composition) detected highly significant, large effects of passage (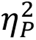 = 0.425), host population age composition (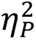 = 0.613), and their interaction (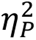 = 0.533) on log-viral load (all *P* < 0.001; Supplementary Table S2). Lineage and passage × lineage terms were also significant.

**Figure 3.**
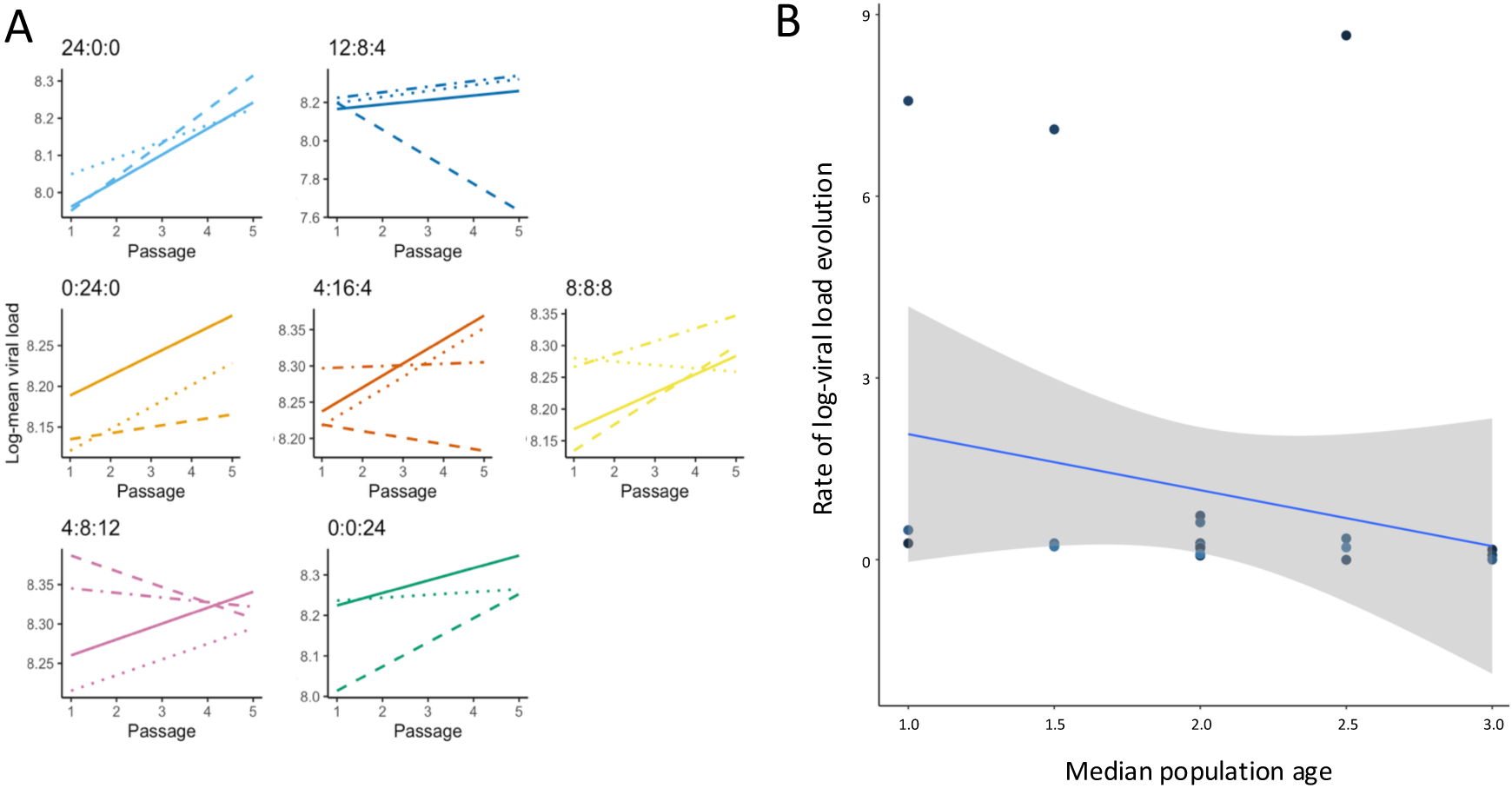
Evolution of viral load and its dependence on host population age composition. (A) Evolution of viral load per infected individual across passages under each host age composition. Different evolutionary lineages are distinguished by line type; host population age compositions are indicated above panels. A GLM detected highly significant, large effects of passage, host age composition, and their interaction on log-viral load (Supplementary Table S2). (B) Dependence of the rate of viral load evolution on the median of the ordinal age categories, Ã. The solid line shows the linear fit (*R*^2^ = 0.045, *F*_1, 24_ = 1.084, *P* = 0.309); the grey band indicates ±1 SD around the regression line.

Consistent with this pathosystem, viral load correlated positively with symptom severity metrics when controlling for Ã: mean severity (*r_p_* = 0.830, 4 d.f., *P* = 0.041), severity at 14 dpi (*r_p_* = 0.952, 4 d.f., *P* = 0.003), AUSIPS (*r_p_* = 0.815, 4 d.f., *P* = 0.048), and PC1 (*r_p_* = 0.965, 4 d.f., *P* = 0.002). These associations support that higher within-host virus accumulation underlies stronger symptom expression in *A. thaliana* - TuMV.

### Host population age composition imposes variable constraints on the rates of evolution for different traits

Rates of phenotypic evolution estimated from time series (ARIMA) revealed distinct dependencies on host demography (Figs. 2C - 2D, S1B - S5B, and 3B). After controlling for independent lineages, the rate of change in PC1 (severity) did not vary detectably with Ã (Fig. 2C; *r_p_* = −0.133, 31 d.f., *P* = 0.462), whereas the rate of change in PC2 (timing) increased with Ã(Fig. 2D; *r_p_* = 0.369, 31 d.f., *P* = 0.034), indicating faster evolution of disease timing in older host populations. By contrast, the rate of viral load evolution showed no measurable association with Ã(Fig. 3B; *r_p_* = −0.219, 22 d.f., *P* = 0.305).

### Specificity of virus adaptation to host population age composition

Cross-infection assays using evolved lineages on hosts populations containing different proportions of individuals at the three developmental stages produced a quantitative infection matrix that was non-nested (WNODF = 7.591, *P* = 1) but strongly modular (*Q* = 0.097, *P* < 0.001) (Fig. 4). TuMV lineages evolved in expansive host populations (Ã = 1 or 1.5) achieved high viral loads only in pre-bolting hosts, consistent with specialization. In contrast, viral lineages evolved in stationary or constrictive populations (Ã = 2 - 3) performed comparably across all developmental stages, suggesting broader host-stage generalism.

**Figure 4.**
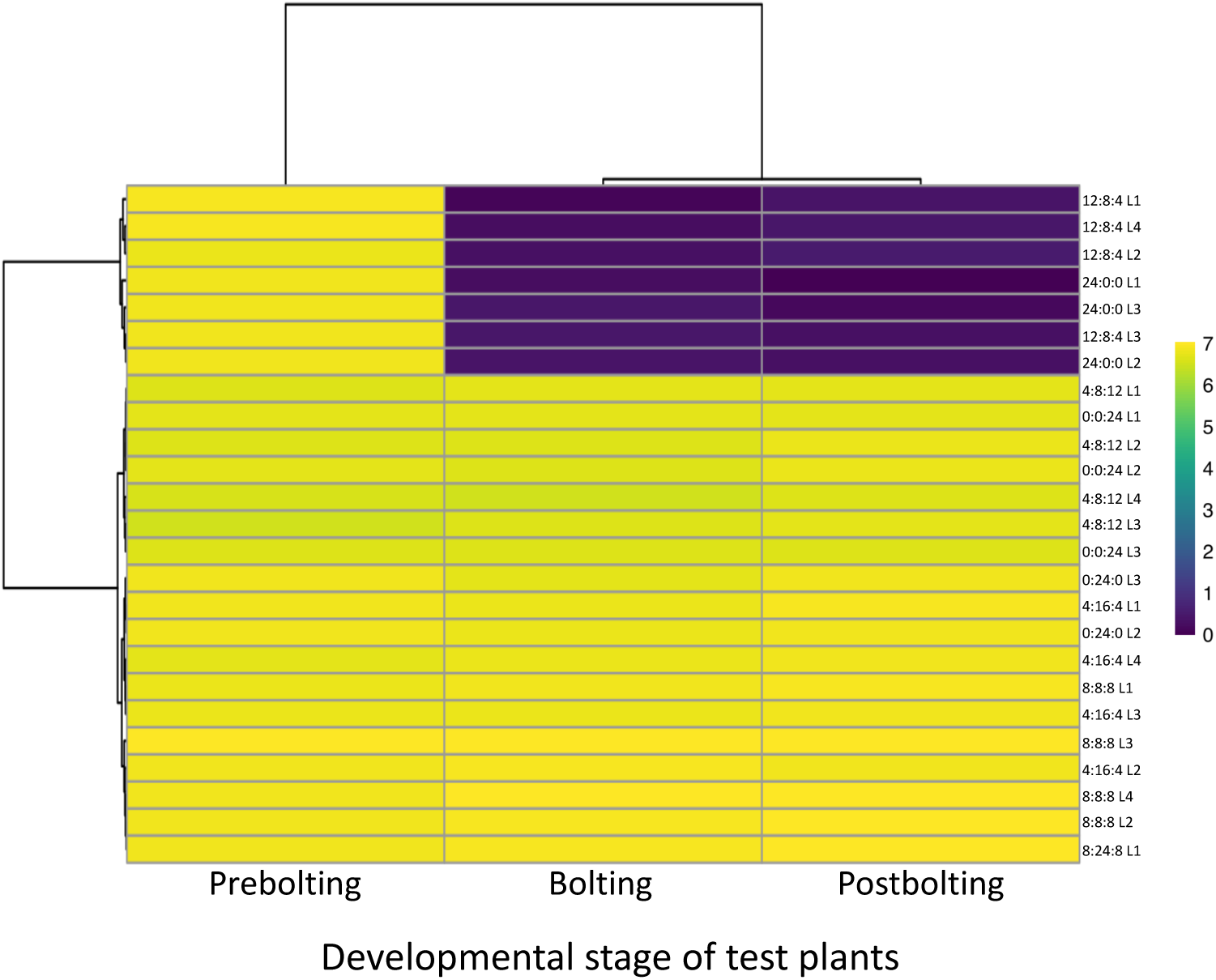
Specificity of adaptation across host developmental stages. Packed quantitative infection matrix summarizing viral load of evolved lineages assayed on hosts at three developmental stages (columns). Rows correspond to lineages evolved under each host population age composition.

### Genetic variability and parallel evolution across demographic treatments

Fig. 5 illustrates the genome-wide changes in genetic variability observed during experimental evolution of TuMV across host populations differing in age structure. Specifically, the figure shows the per site change in heterozygosity (Δ*h*) between passages *t* = 1 and *t* = 5 for all experimental lineages. The distribution of Δ*h* values reveals substantial heterogeneity among genomic sites, with some positions showing sharp increases or decreases in variability, while others remain largely unchanged. These differences are not uniform across demographic treatments: distinct age-structured host populations show unique signatures of heterozygosity change, demonstrating that the host’s demographic composition shapes the evolutionary trajectories of individual viral genomic sites. To statistically evaluate these patterns, we performed an ART ANOVA, which revealed strong and highly significant effects of passage, host population age structure, and their interaction on Δ*h* (Supplementary Fig. S6A; *P* < 0.001 in all cases). Thus, both temporal evolutionary processes and demographic differences in host populations influence viral genetic variability. By contrast, the rate of heterozygosity evolution showed no measurable association with the mean host age Ã (Supplementary Fig. S6B; *P* = 0.120), indicating that although age structure affects patterns of variability, it does not systematically accelerate or decelerate the rate of heterozygosity change across treatments.

**Figure 5.**
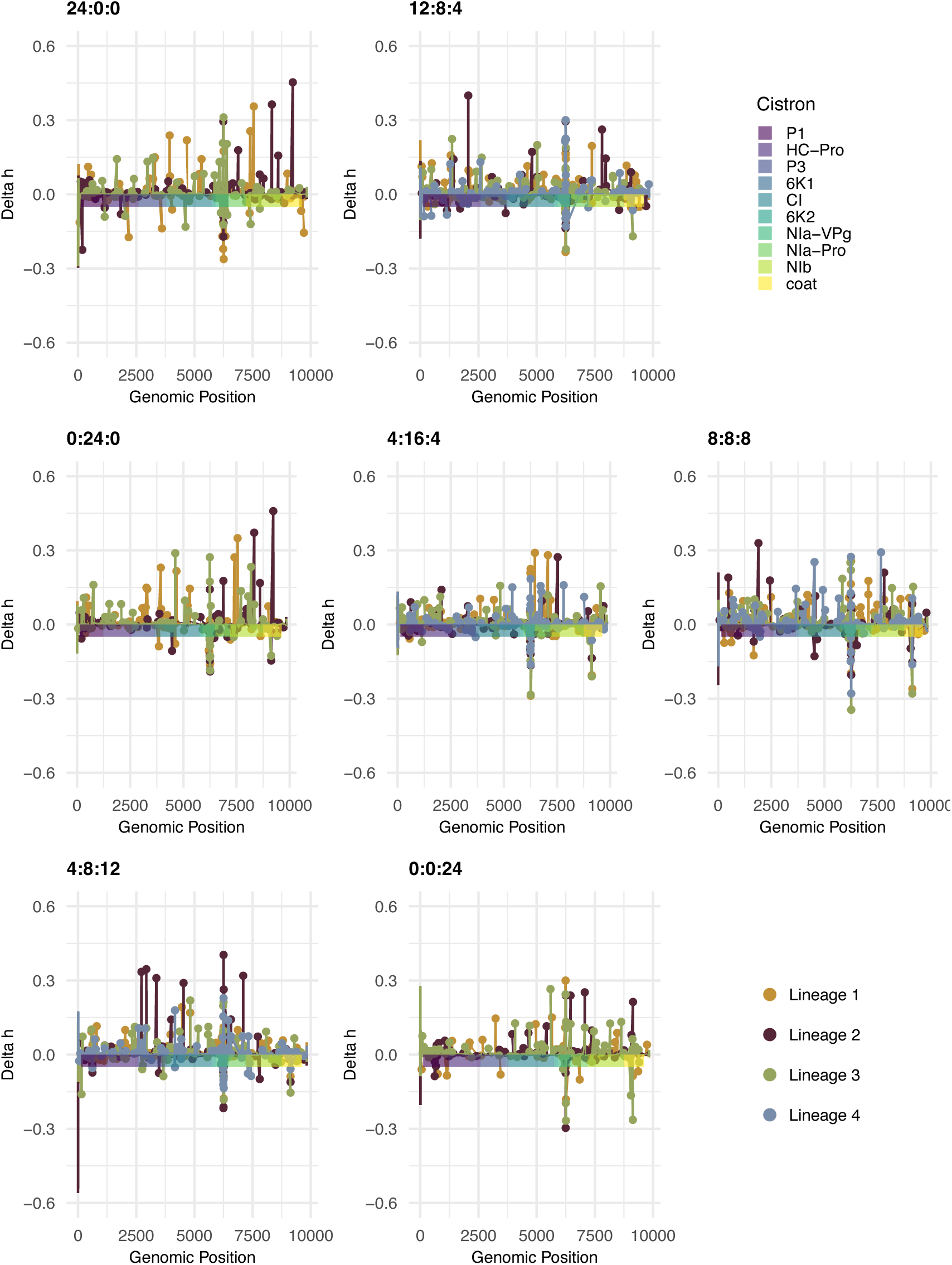
Changes in genetic variability. Differences in per site heterozygosity along TuMV genome between passages 1 and 5 (Δ*h*). Different cistrons are indicated with different colors in the abscissae. Lineages are indicated with colored circles.

To assess whether replicate lineages evolving under the same demographic conditions followed similar genetic trajectories, we performed CMH tests of parallel evolution (Fig. 6). Several demographic treatments exhibited clear parallelism: multiple replicates acquired the same mutation and showed allele-frequency shifts in the same direction, either increases or purging (see Supplementary File S1 for the complete set of cases). The convergent mutation with the largest beneficial effect was 9140A>C (CP/K3004N; basic by polar), observed in all replicates evolved under 0:0:24 (*s* = 0.627 ±0.448; Δ*p* = 0.041). The second-largest beneficial effect corresponded to 3366U>C (synonymous), detected only in 4:8:12 lineages (*s* = 0.460 ±0.567; Δ*p* = 0.048). Remarkably, in 12:8:4, mutations 6243A>G (VPg/N2039D; polar by acid) and 6252C>U (VPg/R2042H; basic by polar) consistently behaved as beneficial across replicates, with large mean selection coefficients per passage (*s* = 0.316 ±0.108 and 0.276 ±0.129, respectively) and substantial increases in frequency (Δ*p* = 0.221 and 0.070), consistent with strong positive selection. Both amino acid replacements occur within a short VPg segment and, taken together, produce a net increase in negative charge. Parallelism was also pervasive among deleterious variants present in the standing genetic variation (Supplementary File S1). The demographic-specific mutation with the largest deleterious effect was 6198G>A (VPg/G2024S; nonpolar by polar), found only in 24:0:0 (*s* = −0.641 ±0.309; Δ*p* = −0.033). In 8:8:8, all lineages shared 6258G>U (VPg/N2044Y; conservative polar) and 9131G>A (synonymous), both strongly deleterious (*s* = −0.994 ±0.341 and −1.094 ±0.067) and undergoing pronounced purifying selection (Δ*p* = −0.194 and −0.124).

**Figure 6.**
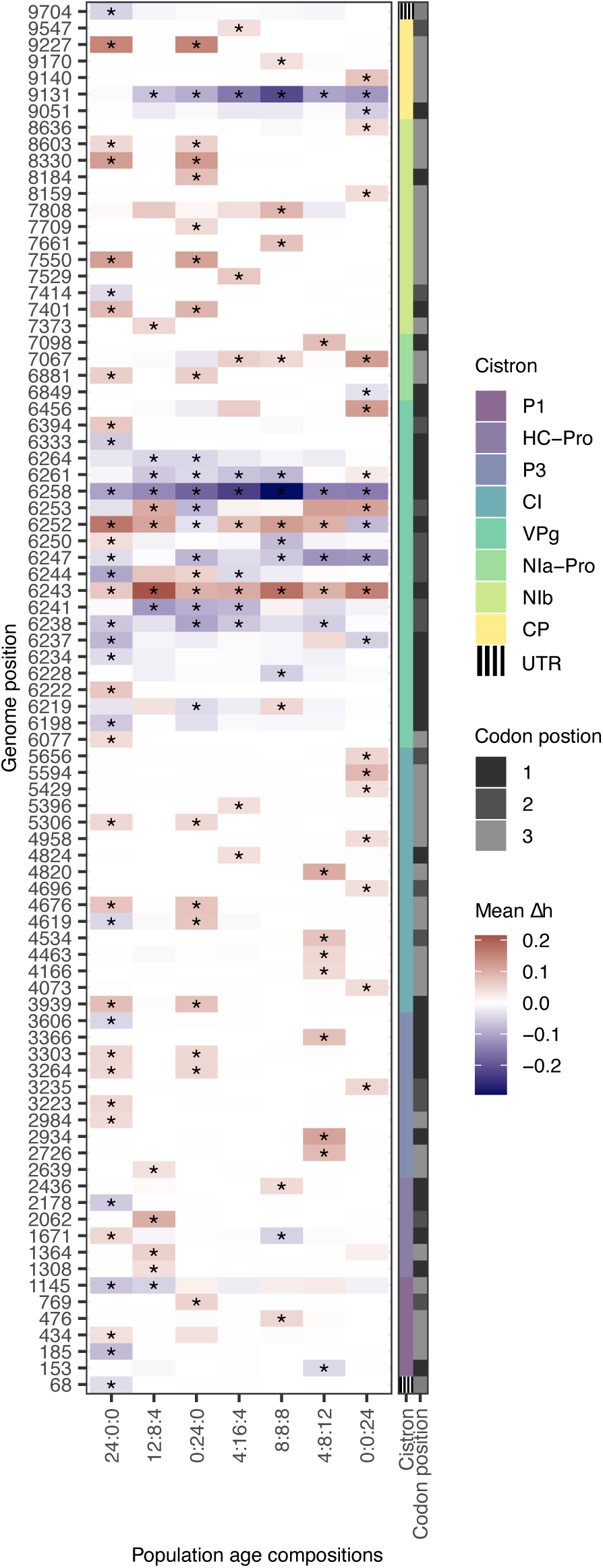
Test of parallelism within and between experimental populations with different demographic compositions.

Several mutations showed treatment-independent deleterious effects (Fig. 6; Supplementary File S1). For example, 6258A>G (VPg/N2044G; polar by nonpolar) was deleterious in all seven treatments, with selection coefficients ranging from −1.069 ±0.134 (Δ*p* = −0.132) in 4:16:4 to −0.510 ±0.502 (Δ*p* = −0.096) in 0:0:24. Mutation 6247A>G (VPg/E2040G; acid by nonpolar) occurred in five demographic conditions (absent in 12:8:4 and 4:16:4), with the strongest negative effect in 4:8:12 (*s* = −0.793 ±0.286; Δ*p* = −0.066) and the weakest in 24:0:0 (*s* = −0.092 ±0.330; Δ*p* = −0.021).

Among synonymous mutations, 9131G>A was deleterious in all demographies except 24:0:0, with selection coefficients ranging from the aforementioned −1.094 in 8:8:8 to −0.410 ±0.384 in 0:24:0 (Δ*p* = −0.054). Likewise, 6261A>G (synonymous) appeared in lineages from five treatments (absent in 24:0:0 and 4:8:12), with *s* ranging from −0.599 ±0.221 (0:24:0; Δ*p* = −0.022) to −0.287 ±0.430 (0:0:24; Δ*p* = −0.056). In contrast, 6243A>G was consistently beneficial across all treatments, with *s* from 0.112 ±0.648 (24:0:0; Δ*p* = 0.226) to 0.456 ±0.094 (0:0:24; Δ*p* = 0.389), indicating a robust advantage irrespective of demography.

A subset of mutations displayed selection coefficients that depended strongly on host age structure, revealing demographic-specific fitness trade-offs. For instance, 6252C>U (VPg/R2042C; basic by nonpolar) was deleterious in constrictive 0:0:24 (*s* = −0.310 ±0.035; Δ*p* = −0.058) and 4:8:12 (*s* = −0.021 ±0.556; Δ*p* = −0.101) demographies, but beneficial in all other, with *s* from 0.009 ±0.518 (0:24:0; Δ*p* = 0.183) to 0.379 ±0.242 (8:8:8; Δ*p* = 0.219).

Two additional notable cases were 6253G>A (VPg/R2042L; basic by nonpolar), which was beneficial in 0:0:24 (*s* = 0.321 ±0.364; Δ*p* = 0.067) and 12:8:4 (*s* = 0.098 ±0.415; Δ*p* = 0.064) but deleterious in 0:24:0 (*s* = −0.491 ±0.408; Δ*p* = −0.045); and 7067C>U (synonymous), beneficial in 0:0:24 (*s* = 0.321 ±0.268; Δ*p* = 0.072) and 8:8:8 (*s* = 0.162 ±0.201; Δ*p* = 0.027) but mildly deleterious in 4:16:4 (*s* = −0.043 ±0.300; Δ*p* = −0.046). These sign reversals indicate an adaptive landscape strongly modulated by host age.

Remarkably, among all the parallel mutations discussed, only VPg/R2042C and VPg/R2042H were *de novo* beneficial mutations, not present in the standing genetic variation present in the first passage, and both affected the same amino acid residue in VPg.

Taken together, Figs. 5 and 6 show that: (*i*) patterns of genetic variability (Δ*h*) are strongly shaped by both evolutionary time and host demographic composition; (*ii*) parallel evolution is common within demographic treatments, with several mutations, including synonymous ones, showing consistent beneficial or deleterious effects across replicates; and (*iii*) across treatments, convergent evolution comprises a mixture of universally beneficial, universally deleterious, and demography-dependent effect mutations, underscoring the complex, multifactorial nature of TuMV adaptation.

## Discussion

### Demography reshapes selection and the virulence-transmission landscape

Our experimental evolution shows that the age composition of host populations imposes distinct and repeatable selective pressures on TuMV, reshaping both the spatial pattern and tempo of genetic change across the viral genome. In particular, we detected strong effects of passage, host age composition, and their interaction on the per-site change in heterozygosity (Δ*h*), whereas the rate of heterozygosity evolution did not scale with the median age Ã (Supplementary Fig. S6). These findings align with classic eco-evolutionary theory linking virulence and transmission to host life history (Anderson & May 1982; Ebert & Herre 1996; Ben-Ami 2019): heterogeneity in host demography shifts the local balance among replication, damage to the host, and onward transmission, thereby moving the location of the virulence-transmission optimum in genotype space. Conceptually, this is consistent with trade-off frameworks for virulence evolution and with the prediction that host life stage can shift the selective gradient experienced by pathogens (for example, via differences in exposure, within-host resources, or immune regulation) (Alizon et al. 2009; Ben-Ami 2019).

Our time-series of disease traits shows that host age structure not only determines whether virulence evolves but which components of virulence respond. Multivariate analyses consolidated five traits into two axes: severity (PC1; driven by AUSIPS, mean severity and severity at 14 dpi) and timing (PC2; dominated by incubation period with opposite loading to AUDPS). Across passages, both axes changed strongly and consistently, but their rates of evolution depended on demography: timing accelerated in older host populations (constrictive), whereas severity did not exhibit a detectable age-dependence. This decoupling implies that age structure can reweight classic virulence components (how fast disease develops *vs* how intense it becomes) thereby shifting epidemiological performance even when severity plateaus. Importantly, viral load (a within-host fitness proxy) increased through passages and correlated positively with severity metrics (and with PC1), linking within-host adaptation to realized symptomatology. Cross-assays further revealed a modular infection matrix: lineages from juvenile-dominated (expansive) host populations specialized on pre-bolting plants, whereas stationary/constrictive populations yielded broader stage generalism. Together, these patterns indicate that host demography redirects the virulence-transmission compromise: juvenile-heavy environments foster rapid gains in stage-specific exploitation (short incubation, high loads in young hosts), while balanced/older environments favor evolvability of timing without necessarily escalating severity. Practically, these results predict distinct epidemic tempos and host-stage targets as fields age, knowledge that can be leveraged through planting schedules or staggered cultivar deployment to slow the ascent of high-virulence phenotypes.

### Phenotypic signatures of virulence under age structure and applied implications

The sign reversals we document (*e.g*., VPg/N2039S, VPg/R2042C or VPg/R2042L) highlight trade-offs: a mutation improving performance in expansive populations can be neutral or deleterious in constrictive populations. This maps naturally onto the idea that host age changes the effective fitness components (replication rate, movement, within-host carrying capacity, and transmission opportunity) that selection sees, shifting the set of optimal genotypes for TuMV in each demographic context. From an applied perspective, these results argue that age structure should be considered a controllable ecological variable when forecasting pathogen evolution or designing durable management (*e.g*., cultivar deployment across developmental windows, staggered planting) and when anticipating cross-stage generalism *vs* stage specialization in evolving lineages.

### Host developmental programs as the ecological driver of selection

From the host side, plants display age-related resistance (ARR) (Kus et al. 2002; Hu & Yang 2019) and ontogenic reprogramming (Carella et al. 2015; Hu & Yang 2019; Melero et al. 2024) of defense, prominently involving salicylic acid (SA) signaling, phase-change regulators (*e.g*., the miR156-SPL module), and differential sensitivity of immune receptors (*e.g*., FLS2) across developmental stages. ARR and ontogenic SA upregulation have been demonstrated in *A. thaliana* and other plants, mechanistically linking development with immune maturation. Recent syntheses highlight that, although general trends exist, ARR is pathosystem-specific and reflects multi-level regulation (RNA silencing, hormones, and transcriptional networks) that can generate stage-dependent selection on viruses, precisely the pattern our data reveal (*e.g*., sign changes in selection coefficients across demographic treatments) (Wilson et al. 2013; Wilson et al. 2017; Melero et al. 2023). Together, our results and this literature suggest that age structure does not merely scale infection risk; it remodels the adaptive landscape encountered by TuMV, shifting which mutations are beneficial, neutral, or deleterious in a demography-dependent manner (*e.g.*, VPg/R2042C beneficial in most treatments but deleterious in constrictive population 0:0:24; Fig. 6).

### Repeatability and context dependence: parallelism within and across demographies

Multiple mutations rose or declined in parallel within the same host demography (*e.g*., beneficials CP/K3004N in 0:0:24 and VPg/N2039 and VPg/R2042H in 12:8:4; deleterious VPg/G2024S in 24:0:0 and VPg/N2044Y in 8:8:8), while several others displayed convergent behavior across different demographic treatments (*e.g*., VPg/N2044G consistently deleterious in all demographies). These outcomes echo extensive evidence that RNA viruses frequently use repeatable genomic routes to adapt to recurring challenges. Parallelism has been observed in experimental systems (*e.g*., vesicular stomatitis virus parallel/convergent mutations across demographic regimes; HIV-1 lineages passaged for ∼1 year) and when viruses adapt to new hosts (*e.g*., Drosophila C virus evolving in 19 species, with stronger parallelism among closely related hosts) (Cuevas et al. 2002; Longdon et al. 2018; Bertels et al. 2019; Bons et al. 2020; Navarro et al. 2022; Ambrós et al. 2024). Clinically and epidemiologically, SARS-CoV-2 provides a recent illustration: independent lineages repeatedly acquire similar spike mutations under host immune pressure and within immunosuppressed patients under antiviral therapy, clear signatures of convergent evolution under shared selection (immune escape, receptor binding) (Roemer et al. 2023). In plant viruses, field and population genomic syntheses emphasize that host ecology and genetic context shape emergence and adaptation, and that repeated solutions can appear in geographically or ecologically distinct settings, patterns mirrored here at laboratory timescales under controlled demography (Elena et al. 2014; Lefeuvre et al. 2019).

Mechanistically, such predictability arises when (*i*) the environment presents a narrow set of functional challenges (Remold 2012), *e.g*., access to translation initiation factors such as eIF4E/eIF(iso)4E for potyviruses, (*ii*) the viral genome contains structurally or functionally constrained hotspots that house large-effect mutations (*e.g*., VPg in potyviruses), and (*iii*) epistasis canalizes trajectories onto a limited number of fit mutational pathways (Sanjuán et al. 2005; Cervera et al. 2016). All three factors have been repeatedly implicated in viral experimental evolution and outbreak analyses. We have showed that some sites are globally selected (*e.g*., VPg/N2044G, beneficial across all treatments), whereas others flip sign depending on demography (*e.g*., VPg/R2042C), underscoring that predictability is conditional on the host context.

### VPg as an evolutionary hinge in potyvirus adaptation

A central outcome of our study is that the majority of recurrent changes map to VPg, and many of them cluster between residues L2003 and N2044. This L2003 - N2044 region has been independently identified as a repeated target of selection in TuMV experimental evolution, with D2037G and R2042H serving as canonical examples of convergent or background-specific solutions in earlier work (Ambrós et al. 2024). The functional salience of this segment is well grounded: VPg mediates key interactions with plant eIF4E/eIF(iso)4E that underlie potyvirus translation, replication, and host specificity, and small amino acid changes within VPg are known to tune or disrupt these contacts (and related interactions), thereby altering fitness in a host- and context-dependent fashion. Beyond protein-factor binding, recent work also shows that evolution in VPg can rewire broader VPg-host proteome interactions, with adaptive substitutions in this very interval modifying the interaction landscape and the balance between growth and defense during infection (Carrasco et al. 2024). The recurrence of VPg hits in our data, especially at positions corresponding to ∼L2003 - N2044, therefore dovetails with independent evidence that this short tract behaves as an evolutionary “hinge” enabling rapid improvement under diverse selective scenarios.

### Synonymous mutations and compositional constraints are visible to selection

A notable feature of our data is parallelism at synonymous sites, including a uniformly beneficial (6243A>G) and deleterious (6261A>G) synonymous changes and demography-dependent synonymous variants (*e.g*., 7067C>U and 9131G>A). That synonymous mutations can be targets of selection is now well established across RNA viruses (Jack et al. 2017; Canale et al. 2018; Le Nouën et al. 2019; Kustin & Stern 2021). Multiple mechanisms can impart strong fitness effects to silent changes: (*i*) Synonymous variants modulate initiation/elongation rates via tRNA availability or codon bias, thereby changing protein yield, stoichiometry across polyproteins, or co-translational folding. Synthetic recoding studies demonstrate that codon and codon deoptimization reproducibly attenuate viruses (poliovirus, influenza A, bacteriophage T7), with measured reductions in mRNA stability and translation efficiency —and importantly, these effects are strictly synonymous. General reviews emphasize that “synonymous is not silent” due to widespread codon bias impacts on expression and fitness (Novella et al. 2004). (*ii*) Silent changes can remodel local or long-range RNA secondary structures required for replication, translation, or packaging; time-series fitness landscapes show many synonymous effects in RNA viruses and their enrichment in structured regions. For positive-sense plant RNA viruses, densely packed genomes impose multiple overlapping constraints (*cis*-acting motifs, frameshifts, RNA-protein interaction sites), magnifying the chance that a synonymous change is functionally visible. (*iii*) In vertebrate systems, elevating CpG/UpA frequencies strongly attenuates viruses via ZAP and OAS3/RNase L pathways; in influenza, engineered CpG-enrichment produces attenuation but robust immunogenicity (Odon et al. 2019). Although plants lack ZAP, plant RNA viruses also show low CpG/UpA and compositional biases shaped by selection and mutation, implying that synonymous changes that shift dinucleotide content or codon usage could influence replication, RNA stability, or recognition by plant defenses (*e.g.*, RNA silencing), with context-dependent fitness effects (González de Prádena et al. 2020; He et al. 2022). These mechanisms provide plausible routes for the recurrent selection we observe at synonymous sites, linked to expression tuning of polyprotein cleavage products (*e.g*., VPg, NIa-Pro, NIb), maintenance of RNA structures, or avoidance/engagement of plant defense layers whose activity varies with host age and tissue ontogeny. The fact that one synonymous variant was beneficial across all demographics (6243A>G), whereas others changed sign with host demography, is consistent with multi-mechanism control: some sites may tune universal functions (*e.g*., robust RNA fold stabilization), while others couple to age-dependent immunity or resource landscapes (*e.g*., shifting tRNA pools or translational capacity across developmental stages).

### Methodological considerations and future directions

We assessed selection from two time points (passages *t* = 1 and *t* = 5) and pooled tissues per passage. However, dense longitudinal sampling (*e.g*., CirSeq-like approaches) would resolve transient polymorphisms and epistatic couplings more finely. Mechanistic validation (reverse genetics) is warranted, particularly for synonymous candidates, to disentangle contributions of RNA structure, codon usage, and dinucleotide content within plant cellular contexts. Finally, measuring age-dependent host traits (hormone profiles, sRNA landscapes, ribosome occupancy, or tRNA pools) would help map specific host changes to viral adaptive solutions [*e.g*., VPg interaction with eIF4E/eIF(iso)4E], refining predictions about when parallelism will be global *vs* demography-specific.

## Conclusions

Our study demonstrates that host age structure is not a passive backdrop but an active driver of viral evolution that alters which mutations are favored, the extent of parallelism, and whether selection is consistent or demography-dependent. The repeated emergence of both nonsynonymous and synonymous changes underscores that multiple layers of constraint — protein function, RNA structure, and translational/ compositional biases— shape predictable paths of adaptation in RNA viruses. By integrating host demography into evolutionary inference, we gain sharper, mechanistically grounded forecasts of how plant viruses will adapt under realistic, age-structured host populations.

## Supplementary material

**Supplementary Figure S1**. Incubation period. (A) Evolution of incubation period under each host population age composition. Different evolutionary lineages are shown by line type. GLM results: significant, mostly large effects of passage (*F*_4, 93.976_ = 51.426, *P* < 0.001, 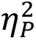 = 0.686), host population age composition (*F*_6, 26.187_ = 41.667, *P* < 0.001, 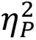 = 0.905), passage × host population age composition (*F*_20, 82.539_ = 2.181, *P* = 0.007, 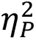 = 0.346), and lineage nested within (passage × host population age composition) (*F*_87, 2118_ = 2.176, *P* < 0.001, 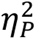 = 0.082). No significant differences were detected among lineages within the same age composition (*F*_27, 168.384_ = 1.979, *P* = 0.896). (B) Rate of evolution of incubation period *vs* the median of the ordinal age categories in each population (partial correlation *r_p_* = 0.448, 31 d.f., *P* = 0.009; controlling for lineage). The solid line shows the linear fit (*R*^2^ = 0.287, *F*_1, 24_ = 9.249, *P* = 0.006); the grey band indicates ±1 SD.

**Supplementary Figure S2.** area under the disease progress stairs (AUDPS). (A) Evolution of AUDPS under each host population age composition. GLM results: passage (*F*_4, 145_ = 39.735, *P* < 0.001, 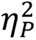 = 0.582), host population age composition (*F*_6, 145_ = 45.693, *P* < 0.001, 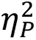 = 0.706), and passage × host population age composition (*F*_20, 145_ = 2.312, *P* = 0.003, 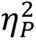 = 0.289) were all significant. (B) Rate of evolution of AUDPS *vs* the median of the ordinal age categories in each population (*r_p_* = −0.048, 31 d.f., *P* = 0.789; controlling for lineage). Solid line: linear fit (*R*^2^ = 0.151, *F*_1, 24_ = 4.083, *P* = 0.055); grey band: ±1 SD.

**Supplementary Figure S3.** Area under symptom intensity progress stairs (AUSIPS). (A) Evolution of AUSIPS under each host age composition. GLM results: passage (*F*_4, 91.651_ = 3.402, *P* = 0.012, 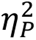 = 0.129), host population age composition (*F*_6, 26.949_ = 6.469, *P* < 0.001, 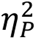 = 0.590), lineages within host population age composition (*F*_27, 85.707_ = 2.082, *P* = 0.006, 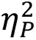 = 0.396), passage × host population age composition (*F*_20, 83.995_ = 3.173, *P* < 0.001, 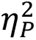 = 0.430), and lineage nested within (passage × host population age composition) (*F*_87, 2117_ = 3.237, *P* < 0.001, 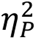 = 0.117)) were all significant. (B) Rate of evolution of AUSIPS *vs* the median of the ordinal age categories in each population (*r_p_* = −0.293, 31 d.f., *P* = 0.098; controlling for lineage). Solid line: linear fit (*R*^2^ = 0.113, *F*_1, 24_ = 2.924, *P* = 0.101); grey band: ±1 SD.

**Supplementary Figure S4.** Mean symptom severity (daily scores). (A) Evolution of mean symptom severity across passages under each host age composition (GLMM on repeated measures). Between-subject effects were significant for passage (*F*_4, 2117_ = 11.837, *P* < 0.001, 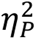 = 0.022), host population age composition (*F*_6, 2117_ = 45.741, *P* < 0.001, 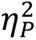 = 0.115), passage × host population age composition (*F*_20, 2117_ = 10.837, *P* < 0.001, 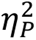 = 0.093), lineages within host population age composition (*F*_27, 2117_ = 7.002, *P* < 0.001, 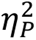 = 0.082), and lineage nested within (passage × host population age composition) (*F*_87, 2117_ = 3.281, *P* < 0.001, 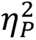 = 0.119). (B) Rate of evolution of mean symptom severity *vs* the median of the ordinal age categories in each population (*r_p_* = 0.050, 31 d.f., *P* = 0.782; controlling for lineage). Solid line: linear fit (*R*^2^ = 0.044, *F*_1, 24_ = 1.062, *P* = 0.313); grey band: ±1 SD.

**Supplementary Figure S5.** Maximum symptom severity at 14 dpi. (A) Evolution of maximum symptom severity at 14 dpi under each host age composition. GLM results: significant, mostly large effects of passage (*F*_4, 92.256_ = 19.676, *P* < 0.001, 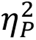 = 0.460), host population age composition (*F*_6, 26.856_ = 3.556, *P* = 0.010, 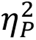 = 0.443), lineages within host population age composition (*F*_27, 85.542_ = 1.868, *P* = 0.016, 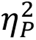 = 0.371), passage × host population age composition (*F*_20, 83.613_ = 2.871, *P* < 0.001, 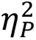 = 0.407), and lineage nested within (passage × host population age composition) (*F*_87, 2118_ = 2.875, *P* < 0.001, 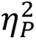 = 0.106). (B) Rate of evolution of maximum symptom severity *vs* the median of the ordinal age categories in each population (*r_p_* = −0.383, 31 d.f., *P* = 0.028; controlling for lineage). Solid line: linear fit (*R*^2^ = 0.148, *F*_1, 24_ = 3.999, *P* = 0.057); grey band: ±1 SD.

**Supplementary Figure S6.** Mean per site nucleotide heterozygosity. (A) Change in mean per site heterozygosity between passages 1 and 5. ART ANOVA results: significant effects of passage (*F*_1, 71404_ = 14.319, *P* < 0.001), host population age composition (*F*_6, 71404_ = 15.037, *P* = 0.010), and passage × host population age composition (*F*_6, 71404_ = 10.859, *P* < 0.001). (B) Rate of evolution of maximum symptom severity *vs* the median of the ordinal age categories in each population (*r_p_* = 0.326, 22 d.f., *P* = 0.120; controlling for lineage). Solid line: linear fit (*R*^2^ = 0.102, *F*_1, 24_ = 2.610, *P* = 0.120); grey band: ±1 SD.

**Supplementary Table S1.** Results of the MANOVA analysis of data shown in Figs. S1A - S5A. Lineages have been added into the model as replicates.

**Supplementary Table S2.** Results of the GLM analysis of log-viral load data shown in Fig. 3A. Lineages have been added into the model as replicates.

**Supplementary File S1.** Results of the CMH test of evolutionary parallelism between lineages evolved under the same demographic conditions (Excel).

## Data and code availability

Raw disease-related phenotypic data and links to R codes used for sequence data analysis and presentation are available at Zenodo repository doi: 10.5281/zenodo.19187176. RNA-seq data were deposited at the European Nucleotide Archive (ENA) at EMBL-EBI under accession number PRJEB97631 (https://www.ebi.ac.uk/ena/browser/view/PRJEB97631).

## Author contributions

S.F.E. conceptualized the project. J.L.C. performed all the experiments. C.T. carried out all the bioinformatic analyses. J.L.C., C.T. and S.F.E. curated the data. C.T. and S.F.E. analyzed the data. S.F.E. wrote the initial manuscript. J.L.C., C.T. and S.F.E. revised the manuscript. S.F.E. secured funding for the project.

## Acknowledgments

We thank Francisca de la Iglesia for excellent technical support, María J. Olmo-Uceda for help with the RNA-seq data analyses and the rest of the EvolSysVir Lab for helpful discussions.

