## Supplementary materials for "Evolution of virulence of a plant RNA virus in age-diverse host populations"

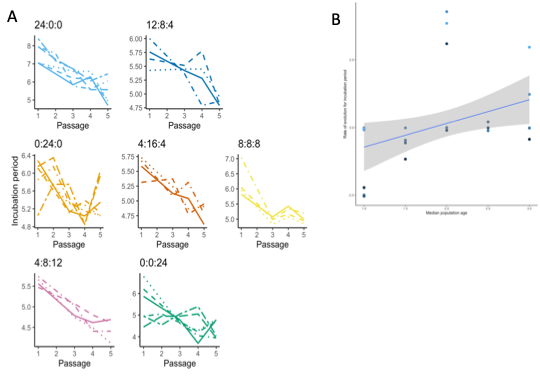

**Figure S1.** Incubation period. (A) Evolution of incubation period under each host population age composition. Different evolutionary lineages are shown by line type. GLM results: significant, mostly large effects of passage (*F*_4, 93.976_ = 51.426, *P* < 0.001, $\eta_{P}^{2}$ = 0.686), host population age composition (*F*_6, 26.187_ = 41.667, *P* < 0.001, $\eta_{P}^{2}$ = 0.905), passage × host population age composition (*F*_20, 82.539_ = 2.181, *P* = 0.007, $\eta_{P}^{2}$ = 0.346), and lineage nested within (passage × host population age composition) (*F*_87, 2118_ = 2.176, *P* < 0.001, $\eta_{P}^{2}$ = 0.082). No significant differences were detected among lineages within the same age composition (*F*_27, 168.384_ = 1.979, *P* = 0.896). (B) Rate of evolution of incubation period *vs* the median of the ordinal age categories in each population (partial correlation *r_p_* = 0.448, 31 d.f., *P* = 0.009; controlling for lineage). The solid line shows the linear fit (*R*^2^ = 0.287, *F*_1, 24_ = 9.249, *P* = 0.006); the grey band indicates ±1 SD.

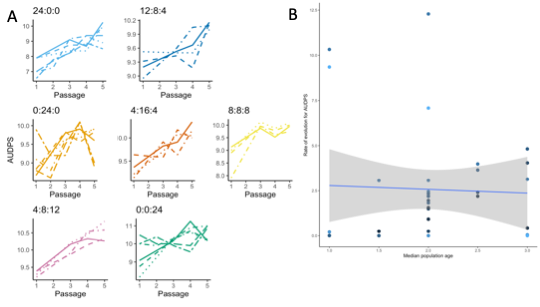

**Figure S2.** Area under the disease progress stairs (AUDPS). (A) Evolution of AUDPS under each host population age composition. GLM results: passage (*F*_4, 145_ = 39.735, *P* < 0.001, $\eta_{P}^{2}$ = 0.582), host population age composition (*F*_6, 145_ = 45.693, *P* < 0.001, $\eta_{P}^{2}$ = 0.706), and passage × host population age composition (*F*_20, 145_ = 2.312, *P* = 0.003, $\eta_{P}^{2}$ = 0.289) were all significant. (B) Rate of evolution of AUDPS *vs* the median of the ordinal age categories in each population (*r_p_* = −0.048, 31 d.f., *P* = 0.789; controlling for lineage). Solid line: linear fit (*R*^2^ = 0.151, *F*_1, 24_ = 4.083, *P* = 0.055); grey band: ±1 SD.

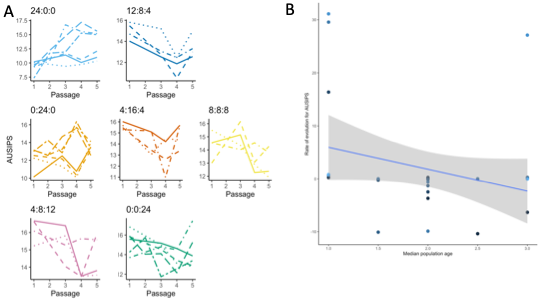

**Figure S3.** Area under symptom intensity progress stairs (AUSIPS). (A) Evolution of AUSIPS under each host age composition. GLM results: passage (*F*_4, 91.651_ = 3.402, *P* = 0.012, $\eta_{P}^{2}$ = 0.129), host population age composition (*F*_6, 26.949_ = 6.469, *P* < 0.001, $\eta_{P}^{2}$ = 0.590), lineages within host population age composition (*F*_27, 85.707_ = 2.082, *P* = 0.006, $\eta_{P}^{2}$ = 0.396), passage × host population age composition (*F*_20, 83.995_ = 3.173, *P* < 0.001, $\eta_{P}^{2}$ = 0.430), and lineage nested within (passage × host population age composition) (*F*_87, 2117_ = 3.237, *P* < 0.001, $\eta_{P}^{2}$ = 0.117)) were all significant. (B) Rate of evolution of AUSIPS *vs* the median of the ordinal age categories in each population (*r_p_* = −0.293, 31 d.f., *P* = 0.098; controlling for lineage). Solid line: linear fit (*R*^2^ = 0.113, *F*_1, 24_ = 2.924, *P* = 0.101); grey band: ±1 SD.

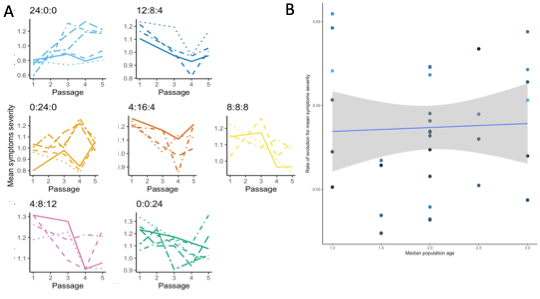

**Figure S4.** Mean symptom severity (daily scores). (A) Evolution of mean symptom severity across passages under each host age composition (GLMM on repeated measures). Between-subject effects were significant for passage (*F*_4, 2117_ = 11.837, *P* < 0.001, $\eta_{P}^{2}$ = 0.022), host population age composition (*F*_6, 2117_ = 45.741, *P* < 0.001, $\eta_{P}^{2}$ = 0.115), passage × host population age composition (*F*_20, 2117_ = 10.837, *P* < 0.001, $\eta_{P}^{2}$ = 0.093), lineages within host population age composition (*F*_27, 2117_ = 7.002, *P* < 0.001, $\eta_{P}^{2}$ = 0.082), and lineage nested within (passage × host population age composition) (*F*_87, 2117_ = 3.281, *P* < 0.001, $\eta_{P}^{2}$ = 0.119). (B) Rate of evolution of mean symptom severity *vs* the median of the ordinal age categories in each population (*r_p_* = 0.050, 31 d.f., *P* = 0.782; controlling for lineage). Solid line: linear fit (*R*^2^ = 0.044, *F*_1, 24_ = 1.062, *P* = 0.313); grey band: ±1 SD.

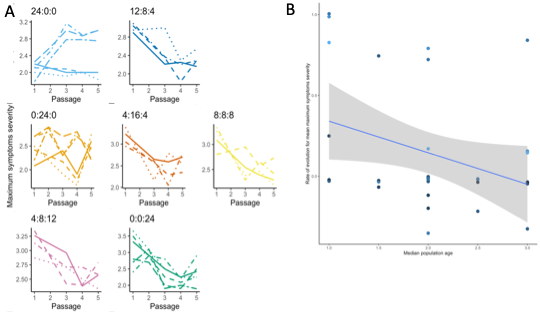

**Figure S5.** Maximum symptom severity at 14 dpi. (A) Evolution of maximum symptom severity at 14 dpi under each host age composition. GLM results: significant, mostly large effects of passage (*F*_4, 92.256_ = 19.676, *P* < 0.001, $\eta_{P}^{2}$ = 0.460), host population age composition (*F*_6, 26.856_ = 3.556, *P* = 0.010, $\eta_{P}^{2}$ = 0.443), lineages within host population age composition (*F*_27, 85.542_ = 1.868, *P* = 0.016, $\eta_{P}^{2}$ = 0.371), passage × host population age composition (*F*_20, 83.613_ = 2.871, *P* < 0.001, $\eta_{P}^{2}$ = 0.407), and lineage nested within (passage × host population age composition) (*F*_87, 2118_ = 2.875, *P* < 0.001, $\eta_{P}^{2}$ = 0.106). (B) Rate of evolution of maximum symptom severity *vs* the median of the ordinal age categories in each population (*r_p_* = −0.383, 31 d.f., *P* = 0.028; controlling for lineage). Solid line: linear fit (*R*^2^ = 0.148, *F*_1, 24_ = 3.999, *P* = 0.057); grey band: ±1 SD.

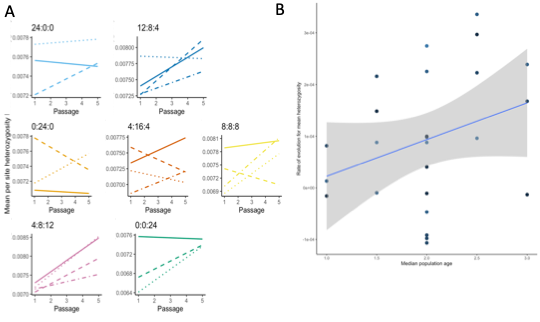

**Figure S6**: Mean per site nucleotide heterozygosity. (A) Change in mean per site heterozygosity between passages 1 and 5. ART ANOVA results: significant effects of passage (*F*_1, 71404_ = 14.319, *P* < 0.001), host population age composition (*F*_6, 71404_ = 15.037, *P* = 0.010), and passage × host population age composition (*F*_6, 71404_ = 10.859, *P* < 0.001). (B) Rate of evolution of maximum symptom severity *vs* the median of the ordinal age categories in each population (*r_p_* = 0.326, 22 d.f., *P* = 0.120; controlling for lineage). Solid line: linear fit (*R*^2^ = 0.102, *F*_1, 24_ = 2.610, *P* = 0.120); grey band: ±1 SD.

| **Table S1.** Results of the MANOVA analysis of data shown in Figs. S1A - S5A. Lineages have been added into the model as replicates. | | | | | | | |
| --- | --- | --- | --- | --- | --- | --- | --- |
| **Effect** | **Wilk’s Λ** | ***F*** | **Hypothesis d.f.** | **Error d.f.** | ***P*** | $\boldsymbol{\eta}_{\boldsymbol{P}}^{\boldsymbol{2}}$ | **Power** |
| Intersection | 0.000 | 13089712.000 | 5 | 110.000 | < 0.001 | 1 | 1.000 |
| Passage | 0.267 | 8.948 | 20 | 365.779 | < 0.001 | 0.281 | 1.000 |
| Host population age composition | 0.206 | 7.139 | 30 | 442.000 | < 0.001 | 0.271 | 1.000 |
| Passage × Host population age composition | 0.262 | 1.708 | 100 | 541.301 | < 0.001 | 0.235 | 1.000 |
| $\eta_{P}^{2}$: magnitude of the effect associated. Values > 0.15 are considered as large effects. | | | | | | | |

| **Table S2.** Results of the GLM analysis of log-viral load data shown in Fig. 3A. Lineages have been added into the model as replicates. | | | | | | |
| --- | --- | --- | --- | --- | --- | --- |
| **Effect** | **Type III SS** | **d.f.** | ***F*** | ***P*** | $\boldsymbol{\eta}_{\boldsymbol{P}}^{\boldsymbol{2}}$ | **Power** |
| Intersection | 9923.905 | 1 | 3279624.107 | < 0.001 | 1.000 | 1.000 |
| Passage | 0.222 | 1 | 73.223 | < 0.001 | 0.425 | 1.000 |
| Host population age composition | 474 | 6 | 26.128 | < 0.001 | 0.613 | 1.000 |
| Passage × Host population age composition | 0.341 | 6 | 18.808 | < 0.001 | 0.533 | 1.000 |
| Lineage(Host age pyramid) | 0.720 | 18 | 13.218 | < 0.001 | 0.706 | 1.000 |
| Passage × Lineage(Host age pyramid) | 0.678 | 18 | 12.455 | < 0.001 | 0.694 | 1.000 |
| Error | 0.300 | 99 |  |  |  |  |
| Total | 10071.979 | 149 |  |  |  |  |
| $\eta_{P}^{2}$: magnitude of the effect associated. Values > 0.15 are considered as large effects. | | | | | | |
